# Phasor Identifier: A Cloud-based Analysis of Phasor-FLIM Data on Python Notebooks

**DOI:** 10.1101/2023.08.30.555392

**Authors:** Mario Bernardi, Francesco Cardarelli

## Abstract

This study aims at creating an accessible notebook tool for the versatile analysis of phasor Fluorescence Lifetime Imaging Microscopy (FLIM) data collected from various samples (e.g. cuvette, cells, tissues) and in various input file formats. The presented strategy facilitates morphological segmentations and diverse mask imports. Results derived from three compelling case studies involving cellular metabolism, nanoscale drug encapsulation (doxorubicin), and the impact of pH and metabolic cleavage on small fluorescent drugs (irinotecan), showcase extensive analysis capabilities. The notebook-centered approach accelerates phasor-FLIM data analysis via external servers, supporting multi-scale research and avoiding the need for GPUs, RAM, and disk space.

## 1 Introduction

Fluorescence Lifetime Imaging Microscopy (FLIM) has revolutionized the field of biological sciences by enabling the investigation and quantification of a multitude of molecular quantities and processes: exemplary applications encompass the measurement of intracellular parameters (e.g. metabolism [1–7], temperature [8–11], viscosity [12– 14]), resolving the physical state of encapsulated fluorescent drugs [15–18]refs, biomedical diagnostics [19–21] and neuroscience research [22–24]. Yet, as recently pointed out by Leonel Malacrida [25], at least two main bottlenecks related to FLIM data analysis still hamper FLIM widespread use, i.e.: *i*) handling the large amounts of data generated can be a significant challenge, particularly for researchers without extensive expertise in data analysis *ii*) post-acquisition data analysis often involves complex mathematical and computational procedures. To analyze FLIM data, lifetime decay curves obtained from microscopy experiments are typically fitted with a variety of mathematical models. While some commercial closed-source software packages (such as those provided by Becker & Hickl, PicoQuant or Leica) and a few open-source options [6–8] were developed to this aim, the analysis of large datasets in a quantitative, easy, fast, and interactive manner remains challenging. In this regard, the phasor approach to FLIM data emerged as a valuable analytical tool. Theoretically introduced by Gregorio Weber already in 1981 [26] and refined shortly after by Jameson and colleagues [27], the phasor approach to lifetime analysis reached widespread dissemination only after a few decades, mainly thanks to its pioneering application to biological samples [28,29]. Worthy of mention, at least two main research trends are stemming out of this approach, i.e. multi-component analysis [30–32] and spectral phasor analysis [33,34]. Also, the wide development of the phasor-FLIM technique elicited efforts for creating new open-source software [35–37] in the attempt to unlock the full potential of FLIM analysis. Here we introduce an innovative strategy that harnesses Google Colaboratory’s (Colab) computational resources for cutting-edge phasor FLIM analysis. Google Colab offers access to powerful computing resources, including GPUs, RAM, and disk space, eliminating the need to invest in costly hardware. Especially if one wants to integrate FLIM designed artificial intelligence tools[38–40] in the notebook. Our goal is to create a user-friendly interface using Google Colab, expanding its use within the biomedical research community. This approach holds the promise of transforming FLIM research by enabling swift, interactive, and quantitative analysis of extensive datasets.

## 2 Materials and Methods

### 2.1 Materials

Liposomal Irinotecan Onivyde® was donated to Scuola Normale Superiore by the Medical Affair Department of Servier Italia S.p.A. One 10-mL vial of sample contains 43 mg Irinotecan anhydrous free base in the form of Irinotecan sucrosofate salt in a pegylated liposomal formulation. The liposomal vesicle is composed of 1,2-distearoyl-sn-glycero-3-phosphocholine (DSPC) 6.81 mg/mL (1:1.6), cholesterol 2.22 mg/mL (1:0.5), and methoxy-terminated polyethylene glycol (MW 2000)-distearoylphosphatidyl ethanolamine (MPEG-2000-DSPE) 0.12 mg/mL (1:0.03). Each mL also contains 2-[4-(2-hydroxyethyl) piperazin-1-yl] ethanesulfonic acid (HEPES) as a buffer 4.05 mg/mL and sodium chloride as an isotonicity reagent 8.42 mg/mL. Irinotecan hydrochloride (powder), purchased from Sigma Aldrich (Milan, Italy), and Onivyde® were both stored at 4°C in compliance with the datasheet. In this study, we evaluated the impact of pH variation on the characteristic lifetime of irinotecan and its metabolite SN-38, purchased from TCI Europe N.V. (Zwijndrecht, Belgium). The pH range studied was from 2.0 to 12.0 and the buffer used was PBS due to its compatibility with living cells and broad buffering capacity. To simplify the methodology, we opted to use PBS rather than more complex buffer mixtures, despite their higher buffering capacity. Nine PBS solutions were prepared with the desired pH, starting from stock solutions of Irinotecan and SN-38 in DMSO. 1-mM solutions in PBS were then prepared for each pH point and the final solutions were stirred to maintain the pH control. For additional insights into the case-study on doxorubicin, readers are referred to Tentori et al.[15]. The Doxoves® used in this study was purchased from FormuMax Scientific (Sunnyvale, CA, USA).

### 2.2 Cell culture

Insulinoma 1E (INS-1E) cells were a kind gift from Professor C. Wollheim from the University of Geneva. These cells were kept in a climate-controlled incubator set to 37°C and 5% CO_2_, where they were grown in RPMI 1640 medium containing 11.1 mmol/L D-glucose, 10 mmol/L HEPES, 2 mmol/L L-Glutamine, 100 U/mL penicillin-streptomycin, 1 mmol/L sodium-pyruvate, and 50 μmol/L β-mercaptoethanol. To conduct lifetime experiments, the cells were allowed to grow until they reached 70% confluence on sterilized microscopy-compatible dishes (IbiTreat μ-Dish 35-mm, Ibidi) for a period of 24-48 hours. Then, the cells were exposed to either Irinotecan or Onivyde® both diluted in complete medium. To serve as a control, the cells were simply refreshed with a fresh batch of complete medium.

### 2.3 FLIM measurements

A drop of approximately 20 μl of Onivyde® was diluted 50x in 980 μl of saline as per intravenous administration protocol. The solution was poured on the glass of a WillCo plate, without any further dilution. For what concerns the free drug, the 1-mM Irinotecan stock solution in DMSO was diluted in different buffers prior to FLIM at a final concentration of ∽10 μM. Irinotecan precipitate and spin-coated liposomes were obtained on the glass of a WillCo plate and black glass bottom 96-well plate respectively, as described above. No aqueous solution was added prior to FLIM to avoid any possible drug re-suspension. FLIM measurements were performed by an Olympus FVMPE-RS microscope coupled with a two-photon Ti:sapphire laser with 80-MHz repetition rate (MaiTai HP, SpectraPhysics) and a FLIM box system for lifetime acquisition (ISS, Urbana Champaign). Onivyde® and Irinotecan were excited at 760 nm and the emission collected by using a 30× planApo silicon immersion objective (NA = 1.0) in the 380-570-nm range. Calibration of the ISS Flimbox system was performed by measuring the known mono-exponential lifetime decay of Fluorescein at pH=11.0 (i.e. 4.0 ns upon excitation at 760 nm, collection range: 570-680 nm). To prepare the calibration sample, a stock of 100 mmol/L Fluorescein solution in EtOH was prepared and diluted in NaOH at pH=11.0 for each calibration measurement. For each measurement, a 512×512 pixels image of FLIM data was collected until 30 frames were acquired. In the context of phasor-FLIM metabolic investigations, living INS-1E cells were observed following 24 hours of cytokine exposure in the standard maintenance conditions. These conditions involved RPMI 1640 medium supplemented with 11.1 mmol/L D-glucose, 10% heat-inactivated FBS, 10 mmol/L HEPES, 2 mmol/L L-Glutamine, 100 U/mL penicillin–streptomycin, 1 mmol/L sodium-pyruvate, and 50 μmol/L β-mercaptoethanol, all maintained at 37°C. We then utilized 2-photon microscopy to capture images of the same cell clusters under both low and high glucose concentrations. Regarding the FLIM measurements related to doxorubicin, an approximately 50 μl droplet of Doxoves® stock solution was carefully applied to the glass surface of a WillCo plate. Notably, no further dilution was carried out. It is important to highlight that this information is distinct from any other FLIM measurements that might have been discussed elsewhere. The actual FLIM process was conducted using a Leica TCS SP5 confocal microscope(Leica Microsystems, Germany). In this setup, a pulsed diode laser operating at a frequency of 40 MHz was employed for excitation, utilizing an excitation wavelength of 470 nm. The emitted light was captured between 520 and 650 nm using a photomultiplier tube, which was linked to a time-correlated single photon counting (TCSPC) card from PicoQuant in Berlin, designated as PicoHarp 300.

### 2.4 Phasor Plot Computations

**Equations (1.a)-(1.b)** describe the computation of the coordinates considering *n* and *ω*, harmonic and angular frequency respectively.

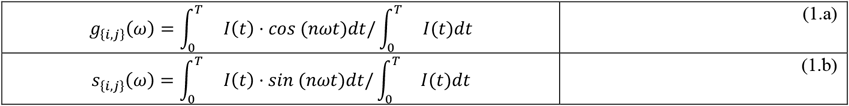

In frequency domain for each pixel one can rely on modulation *m*_*i,j*_ and phase shift *ϕ*_*i,j*_ of the signal as reported in [**Eq. (2.a)-(2.b**)]:

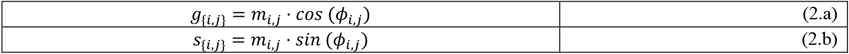

The phasors lie within the semi-circle, also known as “universal semi-circle”, centered at (½,0) with radius ½ and positive x, where the zero lifetime is located at (1,0) and the infinite lifetime at (0,0). Indeed, by taking the Fourier transformation of a measured decay curve, the lifetime can be estimated relying solely on the position of the phasor inside the universal circle. The distribution of phasor points originating from FLIM measurements is found on the universal semi-circle for mono-exponential decays, or within the universal semi-circle for multi-exponential decays. In case of a mono-exponential decay, the intensity can be expressed according to [**Eq. (3.a)**] whereas multi-exponential decay intensity can be expressed according to [**Eq. (3.b)**], where subscripts *f, b*, and *p* indicate different physical states of the fluorescent compound (e.g., Irinotecan in free, membrane-associated and gelated/precipitated forms).

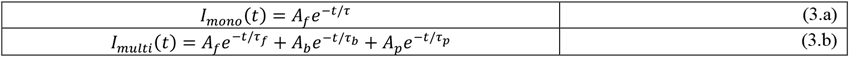

If two molecular species are coexisting in the same pixel, for instance, all the possible weighting combinations of the two molecular species give phasors distributed along a straight line joining the characteristic phasors of the two pure species. In the case of three molecular species, the possible combinations of the system fill a triangle where the vertices correspond again to the characteristic phasors of the pure species. As shown in **Fig. S1** given an experimental phasor that is the combination of two (or more) species, and the phasors of the isolated pure components, a graphical solution can be derived.

## 3 Results and Discussion

### 3.1 Phasor-FLIM Data Analysis: general workflow

In this study, experimental lifetime acquisitions underwent phasor analysis using custom Python 3.6 routines within the *Phasor Identifier* notebook in the Google Colab environment. The input parameters allow specifying the file extension and analysis type, which can be either cumulative or single file. The file name format comprises the date, sample name, and replica number (e.g., **2023_experiment 1_1**). As of today, allowed file formats are: “.R64”, “.ref” and “.ifli”. With the cumulative analysis setting, the code identifies samples by name and stitches images based on replica numbers. For each pixel in the FLIMbox-generated image, the fluorescence decays can be measured in the time-domain (**Fig. 1A**) and mapped onto a “phasor” plot (**Fig. 1B**). The phasor plot has two coordinates, which are the real and imaginary parts of the Fourier transform of the fluorescence lifetime decay. This calculation is performed at the angular repetition frequency of the measurement. Consequently, pixels with similar decay curves exhibit similar coordinates on the phasor plot. **Figure 1B** demonstrates monoexponential decays, indicated by all points lying on the phasor plot. In the case of pixels containing a combination of two or more distinct lifetime decays, they are mapped based on the combination of these contributions. Interpreting the data in three dimensions (**Fig. 1C**) is of utmost importance since there are inevitably regions in the phasor plot that exhibit varying degrees of population. Therefore, proper sampling of these regions is desirable. However, in **Fig. 1C**, the characteristic luminescence signal is projected in 2D for visualization purposes and sampled with 100 bins in both dimensions. It is noteworthy that commonly available open-source codes generate contour plots using elliptical approximation of contour lines. However, this code analyzes the signal through frequency levels, with the bottom level representing the lowest possible level after intensity thresholding, and the top level identified as the 95th percentile. The bottom level is typically discarded to facilitate frequency noise removal. Both the percentile interval and the frequency levels can be fully adjusted and customized.

**Figure 1.**
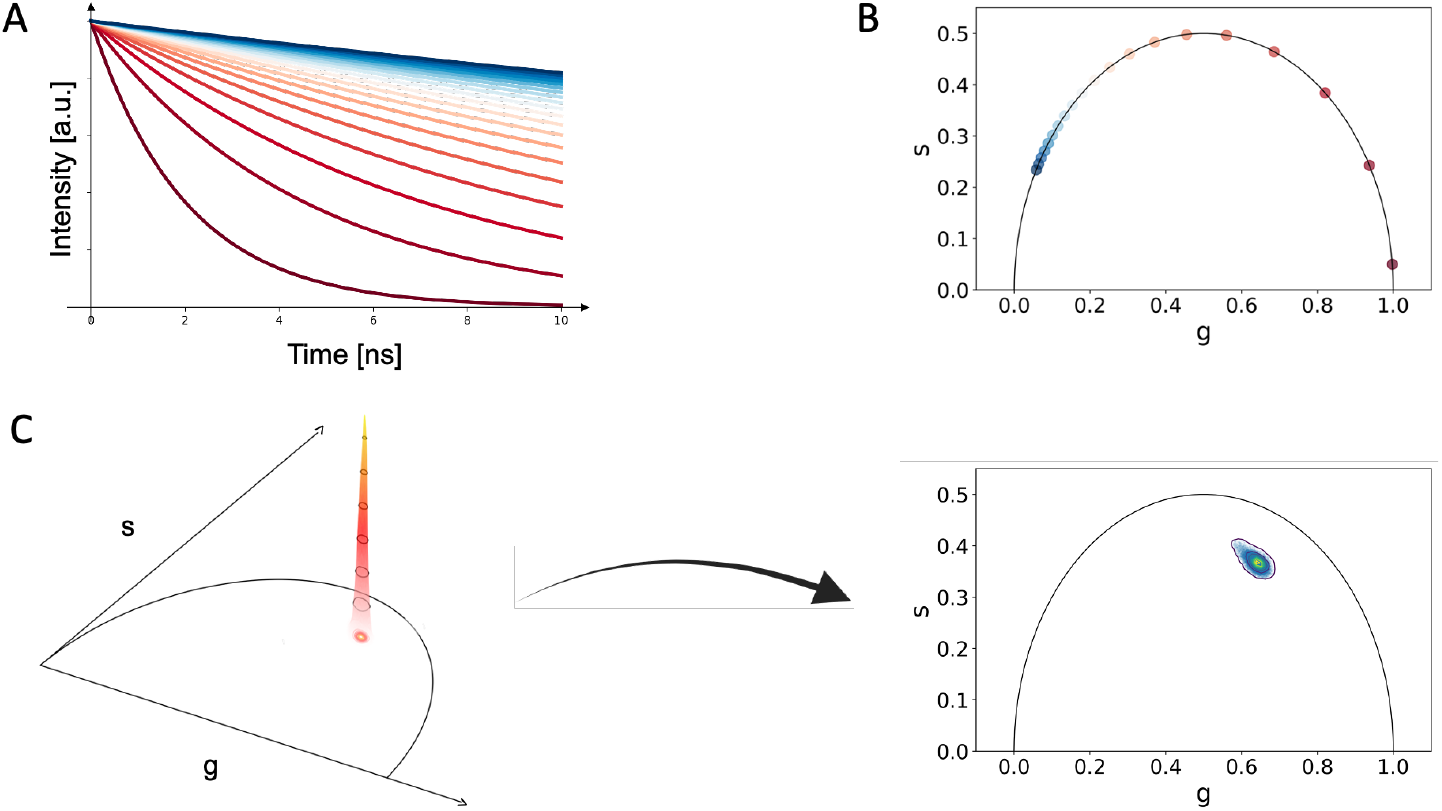
Phasor Analysis of FLIMdata. **A)** A fluorescence decay is measured in the time-domain in each pixel of the image **B)** The fluorescence lifetime decay Fourier transform coordinates are mapped onto a phasor plot, revealing distinct clusters for different decay patterns. **C)** A 2D projection of the phasor plot helps visualize regions with varying population, highlighting the importance of interpreting the data in three dimensions for comprehensive analysis.

The intensity threshold, as shown in **Fig. 2A** on liposomal irinotecan (Onivyde®) samples, can be set through three different approaches: custom threshold value, Otsu threshold, or multi-Otsu threshold. The default option is the multi-Otsu threshold, but setting a custom threshold would be straightforward as intensity values are pre-normalized to 1. Additionally, users have the option to import masks to restrict the analysis to a specific portion of the image, especially when focusing on a particular morphological feature; the code allows for masking cells via the cellpose Python library also. After setting the intensity threshold, the relevant pixels for the analysis are automatically detected. By saving the pixel numbers to be processed, access to the phase and magnitude matrices is restricted to only these relevant pixels, ensuring that the analysis focuses on the desired data points. These points undergo further filtering (**Fig. 2B**), giving users the choice between a Gaussian or median filter. The strength of a median filter refers to the window size used for filtering, where larger values yield stronger smoothing effects. Similarly, the strength of a Gaussian filter is determined by the standard deviation (sigma), with larger sigma values resulting in stronger smoothing. By default, the code sets a median filter strength to 3. The contour analysis function performs a comprehensive analysis of the phasor plot generated FLIM data. It iterates through each frequency level detected in the 3D histogram (**Fig. 2C**) and its corresponding contours, creating a Polygon object for each contour. It then checks if a contour belongs to a previously identified region of interest (ROI) or if it is a new one. A new ROI that satisfies the criteria for a valid contour is added to the perimeters list, which contains the contour data of the identified ROI, as shown in **Fig. 2D**. Similarly, if the contour belongs to an existing ROI, the function updates the corresponding phasor if it is contained within any previously stored phasors; otherwise, it adds the contour as a new phasor as shown in **Fig. 2E**.

**Figure 2.**
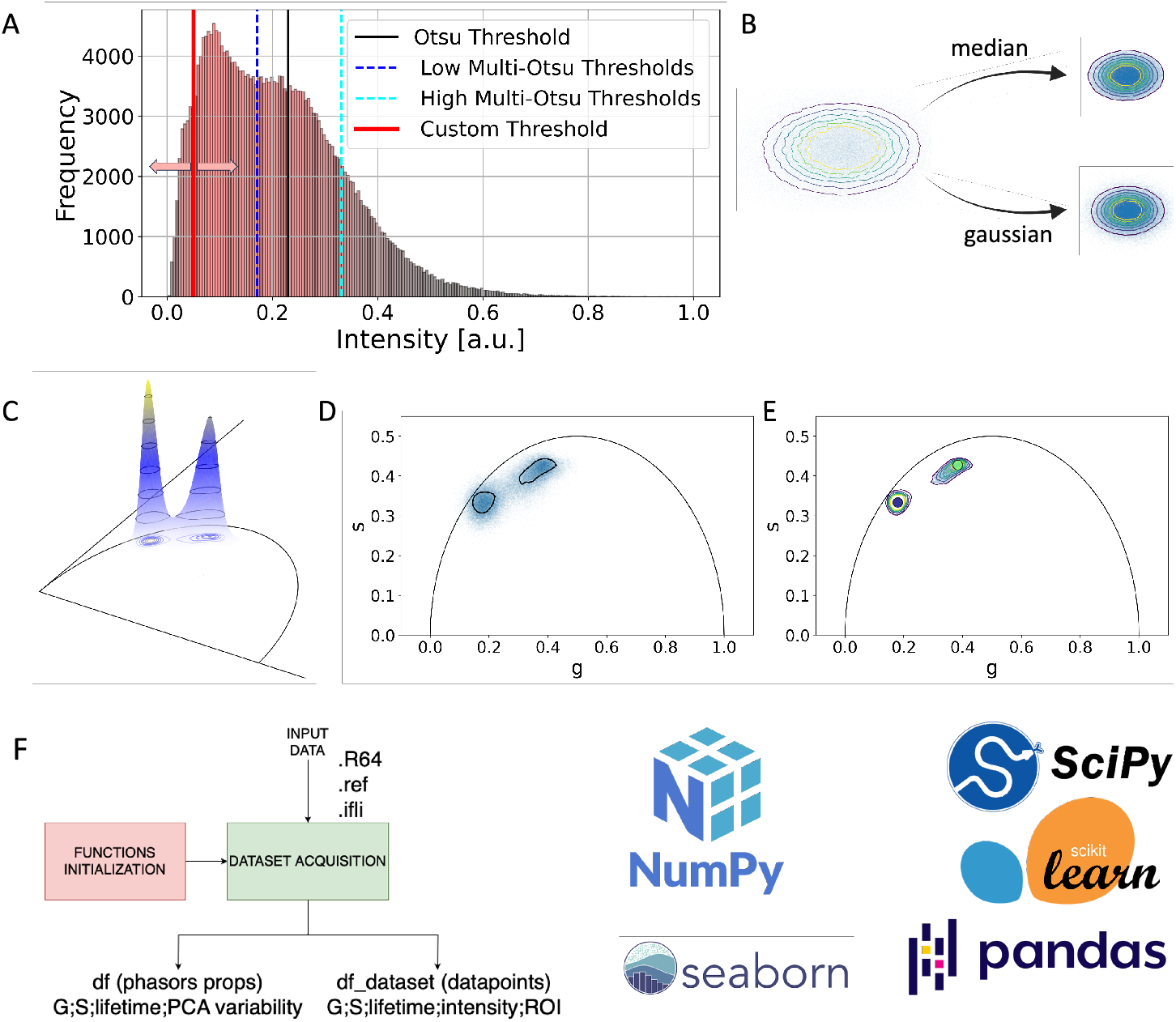
Analysis Process with Filtering and Relevant Pixel Selection. **A)** Automatic detection of relevant pixels based on the set intensity threshold (Custom, Otsu or Multi-Otsu Threshold) **B)** filtering of the relevant pixels using a median or Gaussian filter with user-defined strength **C)** 3D visualization of multi-phasor FLIM signal **D)** Distinct ROIs identification from phasor-FLIM signal in a single sample measurement **E)** Distinct phasors identification from phasor-FLIM signal in a single sample measurement. **F)** The primary objective of the code’s main block is to preprocess and organize data into user-friendly datasets, facilitating subsequent manipulation in subsequent modules.

The valid contour criteria can be set manually, by default new ROIs must contain at least 5000 data points and new phasors must contain at least 500 data points. As a result, the function generates two lists: phasors containing the contour data of identified sub-regions in the phasor plot and perimeters containing the contour data of identified ROIs. These lists provide a comprehensive representation of the regions of interest and sub-regions in the phasor plot, facilitating further analysis and interpretation of the FLIM data. As reported in **Fig. 2F**, the code maintains two distinct dataframes: one, “df_dataset,” storing each datapoint’s data, and another, “df,” documenting phasor properties. These dataframes can be saved as CSV files and accessed beyond the Google Colab notebook for further analysis and visualization.

### 3.2 Time-evolution of phasor-FLIM signals

FLIM is often time used considering the technique sensitivity to chemical, physical, and biological changes. **Fig. 3A** shows these effects considering the phasor position of irinotecan at physiological pH as a reference. A basification of the chemical environment results in longer irinotecan characteristic lifetime signal; irinotecan metabolic enzymatic cleavage into the very potent SN-38 compound leads to shorter lifetime while irinotecan liposomal encapsulation (Onivyde® liposomal formulation) leads to a combination of physical states [16] and interactions with the liposomal membrane that promote longer lifetime values. Interestingly, both metabolic cleavage and encapsulation lead to multi-exponential phasors (hence not lying on the semi-circle in the phasor plot) since the molecule is not found in a single physical state. In fact, as shown in **Fig. 3B**, SN-38 shows an intrinsically multi-exponential phasor as the result of a very dynamic chemical equilibrium. The code allows to follow the evolution of the FLIM signal for each of the three scenarios mentioned above (see **Tab. S1**). However, to keep things straightforward, herein we focus on tracking the lifetime of SN-38 in relation to pH (for SN-38 and irinotecan values see **Fig. S2** and **Tab. S2**). This choice is made because the change in irinotecan lifetime is less apparent, as outlined in **Fig. S2**. Fitting the SN-38 signal in the phasor plot with a linear interpolation (R^2^*≈*0.80) is not very satisfactory. It is rather more convenient to move out of the phasor plot in one variable (i.e., pH) against lifetime space to appreciate a proper non-linear fit (R^2^*≈*0.98). As shown in the box plot of **Fig. 3C**, the code stores all the data points, relying on pandas dataframes, and those can be used to visualize lifetime distributions, in the case as a function of pH. The code stores two different dataframes, one containing the data of each datapoint named df_dataset and one another accounting for the properties of phasors named df. Both can be saved as csv and accessed outside of Google Colab. FLIM is frequently employed due to its sensitivity to changes in chemical, physical, and biological properties. In **Fig. 3A**, these effects are demonstrated through the phasor position of irinotecan under physiological pH, serving as a reference point. Modifying the chemical environment towards basic conditions extends the characteristic lifetime of irinotecan. The metabolization process yielding the potent SN-38 compound shortens the lifetime, while nanoscale encapsulation (into Onivyde® liposomal formulation) introduces diverse physical states and interactions with the liposomal membrane, contributing to longer lifetimes. Intriguingly, both metabolic cleavage and encapsulation lead to multi-exponential phasors, reflecting diverse physical states of the molecule.

**Figure 3.**
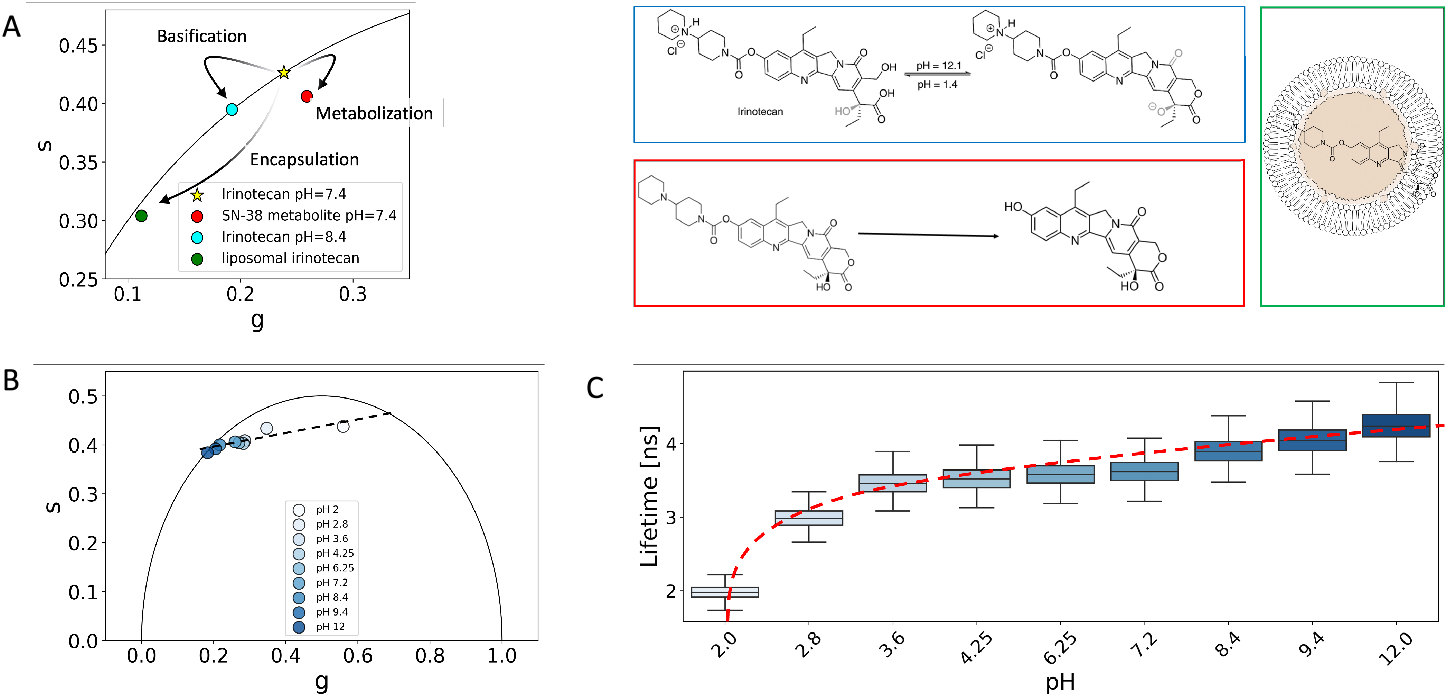
Phasor-FLIM detection and analysis of changes in lifetime. **A)** Phasor-FLIM promptly detects changes in lifetime values stemming from chemical alterations (e.g., basification – depicted in cyan), biological transformations (e.g., metabolic cleavage– highlighted in red), or physical adaptations (e.g., nanoscale encapsulation – in green). In this context, irinotecan (marked with a star) at physiological pH serves as a reference point. **B)** The evolution of SN-38 within the phasor plot is elegantly portrayed, illustrating its response across varying pH levels. **C)** Depicting the pH against lifetime non-linear curve of SN-38, the representation reveals the intricate relationship. Utilizing the seaborn library, the code generates a boxplot derived from dataframes, meticulously stored within the code.

**Figure 3B** reveals that SN-38 inherently exhibits a multi-exponential phasor due to its dynamic chemical equilibrium. Attempting a linear interpolation fit for the SN-38 signal within the phasor plot (R^2^*≈*0.80) proves less satisfactory. This approach doesn’t consider the non-linear relationship between lifetime and phasor plot space, disregarding the influence of pH fluctuations. A more suitable strategy involves transitioning outside of the phasor plot to a space where a variable (e.g., pH) is plotted against lifetime. This facilitates a proper non-linear fit (R^2^*≈*0.98). The box plot in **Fig. 3C**, generated with the seaborn library, showcases data points stored using pandas dataframes.

### 3.3 Morphological and statistical analysis: a glimpse into cellular metabolism

Once all the ROIs have been detected, the code enables morphological analysis on selected ROIs of each sample, chosen through a drop-down menu. The analysis includes computing lifetime and its average over intensity values, along with respective standard deviations. Additionally, it calculates G and S coordinates with corresponding errors, PCA variability ratio, and the number of points sampled to define the phasor. Identifying distinct lifetime regions in a single image is often crucial, so the code facilitates an analysis of intensity, lifetime, and clustering mapping of the sample. Clustering is performed using a Gaussian mixture model[42], which is recommended for phasor-FLIM data. The users have the flexibility to manually specify the number of clusters, with the default set to 2 clusters. **Figure 4 A-B** presents a comparison of INS-1E autofluorescence lifetime signals following incubation with fresh medium (CTRL) and cytokines (CTK). The phasor plot reveals a noticeable shift in the relative lifetime, and there are subpopulations spreading out of the contour plot core. CTRL and CTK study cases were mapped in lifetime, both on the phasor plot and morphologically, leveraging the dataset established by Pugliese et al.[41]. This allows to visually conclude that the CTK treatment has an effect, triggering a shift in both nuclear and cytoplasmic signals.

**Figure 4.**
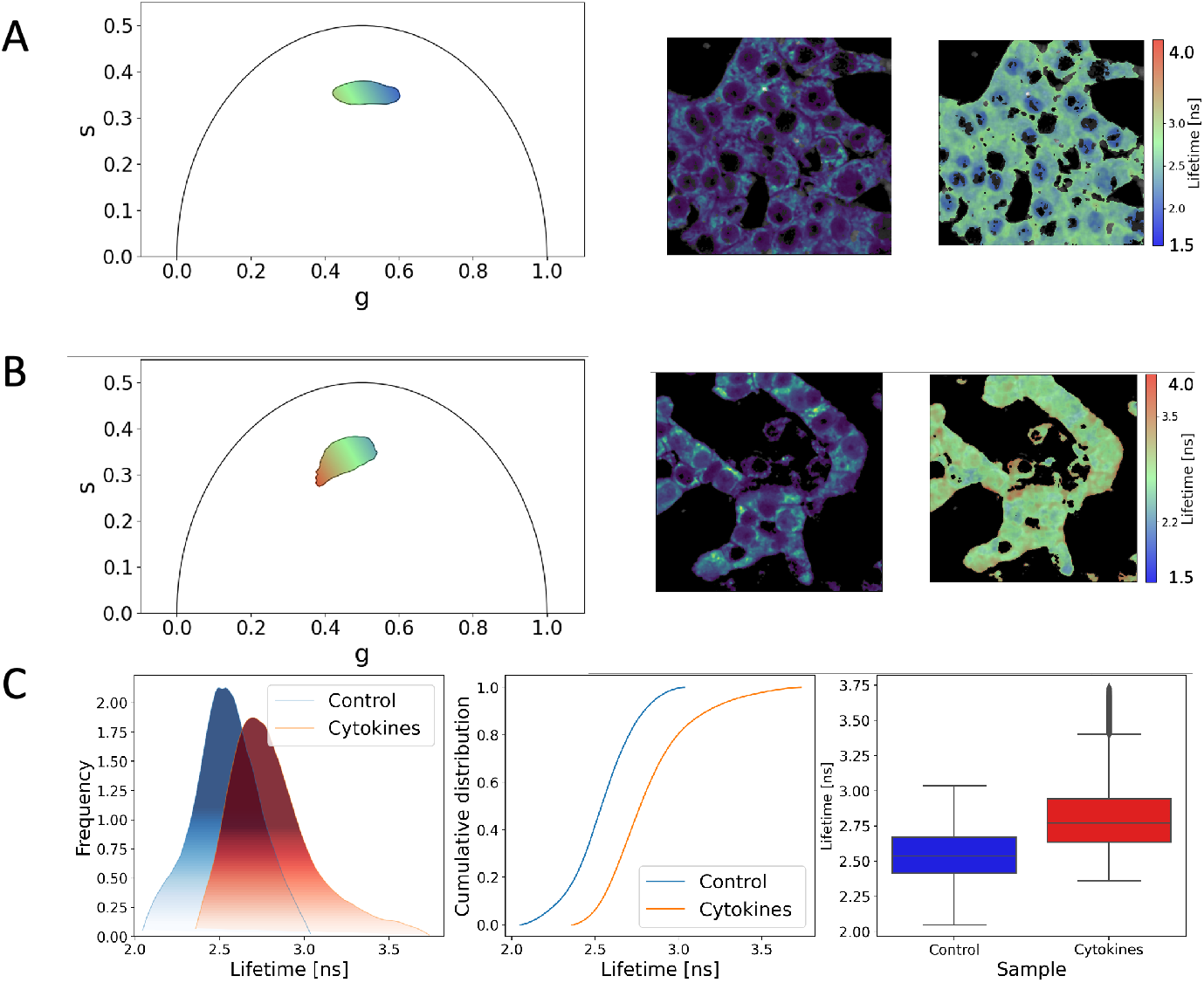
Phasor FLIM Comparative Statistical and Morphological Analysis. **A)** INS-1E autofluorescence lifetime signals after incubation with fresh medium (CTRL) FLIM analysis: in the phasor plot (left panel, color-coded) intensity (middle panel) and lifetime (right panel, color-coded) images **B)** INS-1E autofluorescence lifetime signals after incubation with cytokines (CTK) FLIM analysis: in the phasor plot (left panel, color-coded) intensity (middle panel) and lifetime (right panel, color-coded) images **C)** Assessment of multimodality using non-parametric fitting (Kernel Density Estimate with Gaussian kernel) on the lifetime data from both CTRL and CTK, with visualization through the cumulative distribution function and boxplot to highlight significant differences. (Data extrapolated from the work of Pugliese et al. [41])

Specifically, CTK shows a nuclear lifetime subpopulation with a shorter lifetime, while CTRL shows a cytoplasmic lifetime subpopulation with a longer lifetime. In **Fig. 4C**, we assess this multimodality by sampling the lifetime using non-parametric fitting (Kernel Density Estimate with Gaussian kernel). Subsequently, we conduct two non-parametric statistical tests: Kolmogorov-Smirnoff and Mann Whitney U with Bonferroni correction, to test the null hypothesis that the distributions for CTRL and CTK are the same. The code enables us to perform these statistical tests and has revealed significant differences in terms of percentile distributions, central tendency, skewness, and spread (see **Tab. S3**). Additionally, **Fig. 4C** includes the cumulative distribution function and boxplot, which facilitate the visualization of these statistical differences. In conclusion, the morphological and statistical modules demonstrated in **Fig. 4** provide a robust framework for conducting metabolism assessments, enabling the analysis of cellular metabolism based on cell autofluorescence signals and investigating fluorescent molecule metabolism or interactions within the cellular environment. In this particular scenario, the alteration in the FLIM pattern of cellular autofluorescence can be attributed to the increase of both enzyme-bound NAD(P)H molecules and oxidized lipid species[41].The distribution comparison and analysis are particularly vital for detecting changes in lifetime that may result from interactions or variations in the sample environment, such as temperature fluctuations or pH changes, thereby offering valuable insights into cellular dynamics and metabolic processes.

### 3.4 Intensity and Molar fractions modules: a case study on liposomal doxorubicin

FLIM has been widely used to study multi-exponential signals, where we have at least 2 molecular species or physical states coexisting simultaneously. A classic example of this is the NAD/NADH ratio, which reveals cellular metabolism. However, in this instance, we’re offering a more advanced tool: one that deals with the presence of three molecular species. This scenario arises in the case of the FDA-approved liposomal nanoformulation Doxoves®, which encapsulates doxorubicin, found in three different physical states simultaneously. As Tentori and co-workers demonstrated [15], nanoencapsulated doxorubicin (Doxoves®) exists in three physical states: free in solution, crystallized, and bound to the liposome membrane. Deconvoluting the multi-exponential signal is relatively straightforward, as it involves a linear combination of the three monoexponential phasors on the phasor plot, representing the free, crystal, and bound physical states. However, it is essential to note that this method provides fractional intensity for each physical state: due to differences in molar extinction coefficient (ε) and quantum yield (QY), fractional intensities can be very different from actual molar fractions. To address this point, we have incorporated a molar fraction module where users can input the name, lifetime (in ns), ε, and QY of each physical state they wish to consider.

In **Fig. 5**, we present the potential of this approach through a case study involving the storage of Doxoves® different temperatures (4°C and 37°C) for a duration of 120 days. This study provides valuable insights into the behavior of the three molecular species under varying storage conditions[15]. In **Fig. 5A**, we observe that higher temperature (depicted in red for 37°C) promotes the transformation of doxorubicin’s physical state, leading it towards both the membrane-bound and free-in-solution states. The evolution of the phasor plot at 37°C suggests that doxorubicin crystalline structure within Doxoves® is nearly completely dissolved, and the position of Doxoves® phasor lies along a trajectory resulting from the combined influence of membrane-bound and free-in-solution doxorubicin. This effect becomes evident after 120 days and is strongly reflected in the changes in lifetime (**Fig. 5B**). Indeed, both storage conditions trigger a monoexponential decay in the values of inverse lifetime (1/lifetime), characterized by comparable decay times: τ_4°C_ *≈* 15 days and τ_37°C_ *≈* 19 days. However, the 4°C storage condition maintains a more consistent lifetime signal in the phasor plot, along with a relatively minor increase in lifetime over the 120-day period (∼6%). Conversely, at 37°C under the same storage conditions, there is a noticeable increase in lifetime (∽36%). As detailed in **Tab. 1**, the fractional intensity of crystallized Doxoves® at 37°C experiences a significant decline over 120 days. The Sankey Diagram fluxes in **Fig. 4C** (see also **Tab. 1**) reveal changes in the nanoparticle composition, primarily driven by the dissolution of crystallized doxorubicin. This results in a new molar composition of the liposomal drug (refer to **Tab. S4** for doxorubicin QY and ε values in the relevant physical states).

**Table 1.** Fractional Intensities and molar fractions of Doxoves® in different storage conditions. All results are obtained from 6 replica and the error on both fractional intensity and molar fraction was systematically <0.005. (Median filter extent = 3).

| Sample | Intensity fraction crystal | Intensity fraction free | Intensity fraction bound | Molar fraction crystal | Molar fraction free | Molar fraction bound |
| --- | --- | --- | --- | --- | --- | --- |
| Doxoves® | 0.459 | 0.125 | 0.416 | 0.988 | 0.007 | 0.005 |
| Doxoves® 4°C<br>120 days | 0.373 | 0.163 | 0.464 | 0.982 | 0.011 | 0.069 |
| Doxoves® 37°C<br>120 days | 0.024 | 0.227 | 0.749 | 0.701 | 0.173 | 0.126 |

**Figure 5.**
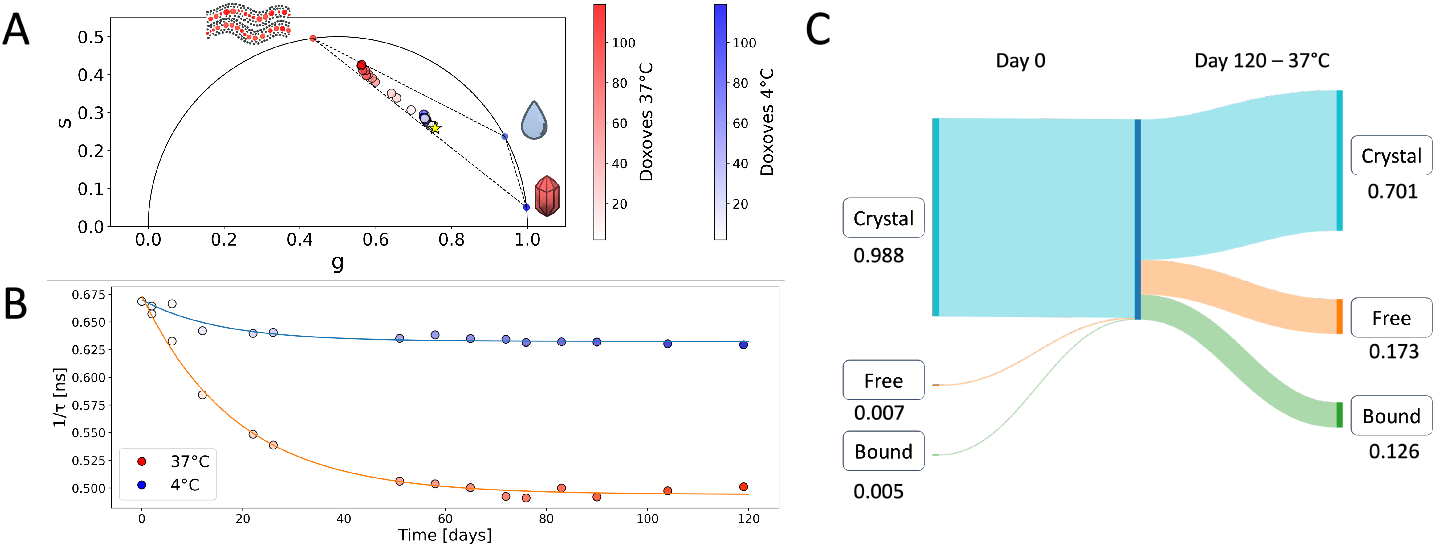
FLIM analysis of storage-induced changes in nano encapsulated drugs: liposomal doxorubicin case study. **A)** The phasor-plot showcases the lifetimes of liposomal doxorubicin (Doxoves®) within the manufacturer’s solution. Monitoring occurred over a 0–120-day period under two distinct storage conditions: 4°C (depicted in the blue palette) and 37°C (depicted in the red palette). The star denotes the initial FLIM signal of Doxoves® on day 0. **B)** The progression of 1/Lifetime values over time unveils a consistent monoexponential decay pattern in both storage conditions, despite the significant variations between the two. **C)** Sankey Diagram of the molar compositional changes of Doxoves® nanoparticles after 120 days of storage at 37°C.

## Supporting information

Comprehensive overviews of phasor and lifetime measurements

## Acknowledgments

The authors kindly acknowledge Dr. Licia Anna Pugliese and Dr. Annalisa Carretta for generously providing part of the lifetime data analysed in this study. The Sankey Diagram of Figure 5 was realized in the open environment provided at https://sankeymatic.com/build/. Phasor positions and lifetime values shown in this paper are in keeping, and were systematically compared with, the outcomes of SimFCS. The authors acknowledge the support of the European Union by the Next Generation EU project ECS00000017 ‘Ecosistema dell’Innovazione’ Tuscany Health Ecosystem (THE, PNRR, Spoke 4: Nanotechnologies for diagnosis and therapy). The work is supported in part by the European Research Council (ERC) under the European Union’s Horizon 2020 research and innovation programme (grant agreement No 866127, project CAPTUR3D).

## Disclosures

The authors declare no conflict of interest.

## Code and Data Availability

The archived version of the code described in this manuscript can be freely accessed through GitHub [https://github.com/Mariochem92/PhasorIdentifier; DOI: 10.5281/zenodo.8282839]. Data are available from the authors upon request.

## Supplemental Document

Schematic representation of the graphical approach for assessing intensity fractions of coexisting species; comprehensive overview of the phasor measurements; comprehensive overview of the lifetime measurements of SN-38 and irinotecan in PBS over the wide range of pH values under study, from 2 to 12; comparison of the fluorescence lifetime of irinotecan and SN-38 at pH values between 2 and 12; lifetime distribution data statistical comparison of INS1E cells (control) and Cytokines; Quantum Yield and molar extinction coefficient of doxorubicin in distinct physical states: crystal, membrane-bound, and free in solution.

## Notes

https://github.com/Mariochem92/PhasorIdentifier

### Competing Interest Statement

The authors have declared no competing interest.

## References

1. R. Scholz, R. G. Thurman, J. R. Williamson, B. Chance, and T. Bücher, “Flavin and pyridine nucleotide oxidation-reduction changes in perfused rat liver. I. Anoxia and subcellular localization of fluorescent flavoproteins,” J Biol Chem 244(9), 2317–2324 (1969).

2. J. R. Lakowicz, H. Szmacinski, K. Nowaczyk, and M. L. Johnson, “Fluorescence lifetime imaging of free and protein-bound NADH.,” Proc Natl Acad Sci U S A 89(4), 1271–1275 (1992).

3. C. Stringari, A. Cinquin, O. Cinquin, M. A. Digman, P. J. Donovan, and E. Gratton, “Phasor approach to fluorescence lifetime microscopy distinguishes different metabolic states of germ cells in a live tissue,” Proceedings of the National Academy of Sciences 108(33), 13582–13587 (2011).

4. C. Stringari, J. L. Nourse, L. A. Flanagan, and E. Gratton, “Phasor Fluorescence Lifetime Microscopy of Free and Protein-Bound NADH Reveals Neural Stem Cell Differentiation Potential,” PLOS ONE 7(11), e48014 (2012).

5. G. Ferri, M. Tesi, F. Massarelli, L. Marselli, P. Marchetti, and F. Cardarelli, “Metabolic response of Insulinoma 1E cells to glucose stimulation studied by fluorescence lifetime imaging,” FASEB BioAdvances 2(7), 409–418 (2020).

6. F. Azzarello, L. Pesce, V. De Lorenzi, G. Ferri, M. Tesi, S. Del Guerra, P. Marchetti, and F. Cardarelli, “Single-cell imaging of α and β cell metabolic response to glucose in living human Langerhans islets,” Commun Biol 5(1), 1–10 (2022).

7. T. S. Blacker, Z. F. Mann, J. E. Gale, M. Ziegler, A. J. Bain, G. Szabadkai, and M. R. Duchen, “Separating NADH and NADPH fluorescence in live cells and tissues using FLIM,” Nat Commun 5(1), 3936 (2014).

8. K. Okabe, N. Inada, C. Gota, Y. Harada, T. Funatsu, and S. Uchiyama, “Intracellular temperature mapping with a fluorescent polymeric thermometer and fluorescence lifetime imaging microscopy,” Nat Commun 3(1), 705 (2012).

9. S. Uchiyama, C. Gota, T. Tsuji, and N. Inada, “Intracellular temperature measurements with fluorescent polymeric thermometers,” Chem. Commun. 53(80), 10976–10992 (2017).

10. M. Suzuki and T. Plakhotnik, “The challenge of intracellular temperature,” Biophys Rev 12(2), 593–600 (2020).

11. A. Murakami, K. Nagao, R. Sakaguchi, K. Kida, Y. Hara, Y. Mori, K. Okabe, Y. Harada, and M. Umeda, “Cell-autonomous control of intracellular temperature by unsaturation of phospholipid acyl chains,” Cell Rep 38(11), 110487 (2022).

12. E. O. Puchkov, “Intracellular viscosity: Methods of measurement and role in metabolism,” Biochem. Moscow Suppl. Ser. A 7(4), 270–279 (2013).

13. T. Liu, X. Liu, D. R. Spring, X. Qian, J. Cui, and Z. Xu, “Quantitatively Mapping Cellular Viscosity with Detailed Organelle Information via a Designed PET Fluorescent Probe,” Sci Rep 4(1), 5418 (2014).

14. M. K. Kuimova, S. W. Botchway, A. W. Parker, M. Balaz, H. A. Collins, H. L. Anderson, K. Suhling, and P. R. Ogilby, “Imaging intracellular viscosity of a single cell during photoinduced cell death,” Nature Chem 1(1), 69–73 (2009).

15. P. Tentori, G. Signore, A. Camposeo, A. Carretta, G. Ferri, P. Pingue, S. Luin, D. Pozzi, E. Gratton, F. Beltram, G. Caracciolo, and F. Cardarelli, “Fluorescence lifetime microscopy unveils the supramolecular organization of liposomal Doxorubicin,” Nanoscale 14(25), 8901–8905 (2022).

16. Bernardi, Mario, Signore Giovanni Moscardini, Aldo, Pugliese, Licia Anna, Pesce, Luca, Beltram, Fabio, and Cardarelli, Francesco, “Fluorescence lifetime nanoscopy of liposomal irinotecan Onivyde®: from manufacturing to intracellular processing,” ACS Appl. Bio Mater. (2023).

17. M. Bernardi and F. Cardarelli, “Nanoscopy on drug-encapsulating nanosystems by phasor-based fluorescence lifetime analysis,” Biophysical Journal 122(3), 278a (2023).

18. F. Cardarelli, F. Beltram, P. M. Tentori, G. Caracciolo, and D. Pozzi, “Determination of the Supramolecular Organization of Encapsulated Molecules by Luminescence Lifetime Analysis,” U.S. patent WO2022097108 (A1) (May 12, 2022).

19. L. Marcu, “Fluorescence Lifetime Techniques in Medical Applications,” Ann Biomed Eng 40(2), 304–331 (2012).

20. A. Alfonso-Garcia, S. Cevallos, J.-Y. Lee, C. Li, J. Bec, A. Baumler, and L. Marcu, “Intraluminal fluorescence lifetime imaging (FLIm) as a diagnostic tool for gastrointestinal disease,” in Biophotonics Congress 2021 (2021), Paper DM3A.4 (Optica Publishing Group, 2021), p. DM3A.4.

21. C. Maibohm, F. Silva, E. Figueiras, P. T. Guerreiro, M. Brito, R. Romero, H. Crespo, and J. B. Nieder, “SyncRGB-FLIM: synchronous fluorescence imaging of red, green and blue dyes enabled by ultra-broadband few-cycle laser excitation and fluorescence lifetime detection,” Biomed. Opt. Express, BOE 10(4), 1891–1904 (2019).

22. R. Yasuda, “3 - Principle and Application of Fluorescence Lifetime Imaging for Neuroscience: Monitoring Biochemical Signaling in Single Synapses Using Fluorescence Lifetime Imaging,” in Neurophotonics and Biomedical Spectroscopy, R. R. Alfano and L. Shi, eds., Nanophotonics (Elsevier, 2019), pp. 53–64.

23. A. J. Bowman, C. Huang, M. J. Schnitzer, and M. A. Kasevich, “Wide-field fluorescence lifetime imaging of neuron spiking and sub-threshold activity in vivo,” Science 380(6651), 1270–1275 (2023).

24. D. J. Liput, T. A. Nguyen, S. M. Augustin, J. O. Lee, and S. S. Vogel, “A Guide to Fluorescence Lifetime Microscopy and Förster’s Resonance Energy Transfer in Neuroscience,” Curr Protoc Neurosci 94(1), e108 (2020).

25. L. Malacrida, “Phasor plots and the future of spectral and lifetime imaging,” Nat Methods 20(7), 965–967 (2023).

26. G. Weber, “Resolution of the fluorescence lifetimes in a heterogeneous system by phase and modulation measurements,” J. Phys. Chem. 85(8), 949–953 (1981).

27. D. M. Jameson, E. Gratton, and R. D. Hall, “The Measurement and Analysis of Heterogeneous Emissions by Multifrequency Phase and Modulation Fluorometry,” Applied Spectroscopy Reviews 20(1), 55–106 (1984).

28. M. A. Digman, V. R. Caiolfa, M. Zamai, and E. Gratton, “The Phasor Approach to Fluorescence Lifetime Imaging Analysis,” Biophysical Journal 94(2), L14–L16 (2008).

29. L. Malacrida, S. Ranjit, D. Jameson, and E. Gratton, “The Phasor Plot: A Universal Circle to Advance Fluorescence Lifetime Analysis and Interpretation,” Annual Review of Biophysics 50, 575–593 (2021).

30. A. Vallmitjana, B. Torrado, A. Dvornikov, S. Ranjit, and E. Gratton, “Blind Resolution of Lifetime Components in Individual Pixels of Fluorescence Lifetime Images Using the Phasor Approach,” J Phys Chem B 124(45), 10126–10137 (2020).

31. A. Vallmitjana, A. Dvornikov, B. Torrado, D. M. Jameson, S. Ranjit, and E. Gratton, “Resolution of 4 components in the same pixel in FLIM images using the phasor approach,” Methods Appl Fluoresc 8(3), 035001 (2020).

32. S. Jeong, D. A. Greenfield, M. Hermsmeier, A. Yamamoto, X. Chen, K. F. Chan, and C. L. Evans, “Time-resolved fluorescence microscopy with phasor analysis for visualizing multicomponent topical drug distribution within human skin,” Sci Rep 10(1), 5360 (2020).

33. G. O. S. Williams, E. Williams, N. Finlayson, A. T. Erdogan, Q. Wang, S. Fernandes, A. R. Akram, K. Dhaliwal, R. K. Henderson, J. M. Girkin, and M. Bradley, “Full spectrum fluorescence lifetime imaging with 0.5 nm spectral and 50 ps temporal resolution,” Nat Commun 12(1), 6616 (2021).

34. L. Scipioni, A. Rossetta, G. Tedeschi, and E. Gratton, “Phasor S-FLIM: a new paradigm for fast and robust spectral fluorescence lifetime imaging,” Nat Methods 18(5), 542–550 (2021).

35. S. Ranjit, L. Malacrida, D. M. Jameson, and E. Gratton, “Fit-free analysis of fluorescence lifetime imaging data using the phasor approach,” Nat Protoc 13(9), 1979–2004 (2018).

36. D. Gottlieb, B. Asadipour, T. P. L. Ung, and C. Stringari, “FLUTE: a Python GUI for interactive phasor analysis of FLIM data,” 2023.03.31.534529 (2023).

37. W. Schrimpf, A. Barth, J. Hendrix, and D. C. Lamb, “PAM: A Framework for Integrated Analysis of Imaging, Single-Molecule, and Ensemble Fluorescence Data,” Biophysical Journal 114(7), 1518–1528 (2018).

38. A. Rossetta, “The bright future of fluorescence lifetime analysis,” in Reporters, Markers, Dyes, Nanoparticles, and Molecular Probes for Biomedical Applications XIV (SPIE, 2023), 12398, pp. 19–22.

39. L. Héliot and A. Leray, “Simple phasor-based deep neural network for fluorescence lifetime imaging microscopy,” Sci Rep 11(1), 23858 (2021).

40. J. T. Smith, R. Yao, N. Sinsuebphon, A. Rudkouskaya, N. Un, J. Mazurkiewicz, M. Barroso, P. Yan, and X. Intes, “Fast fit-free analysis of fluorescence lifetime imaging via deep learning,” Proceedings of the National Academy of Sciences 116(48), 24019–24030 (2019).

41. L. A. Pugliese, V. De Lorenzi, M. Bernardi, S. Ghignoli, M. Tesi, P. Marchetti, L. Pesce, and F. Cardarelli, “Unveiling nanoscale optical signatures of cytokine-induced β-cell dysfunction,” Sci Rep 13(1), 13342 (2023).

42. A. Vallmitjana, B. Torrado, and E. Gratton, “Phasor-based image segmentation: machine learning clustering techniques,” Biomed. Opt. Express, BOE 12(6), 3410–3422 (2021).

