## Supplementary material for "Phasor Identifier: A Cloud-based Analysis of Phasor-FLIM Data on Python Notebooks": Comprehensive overviews of phasor and lifetime measurements

### Supplementary Information

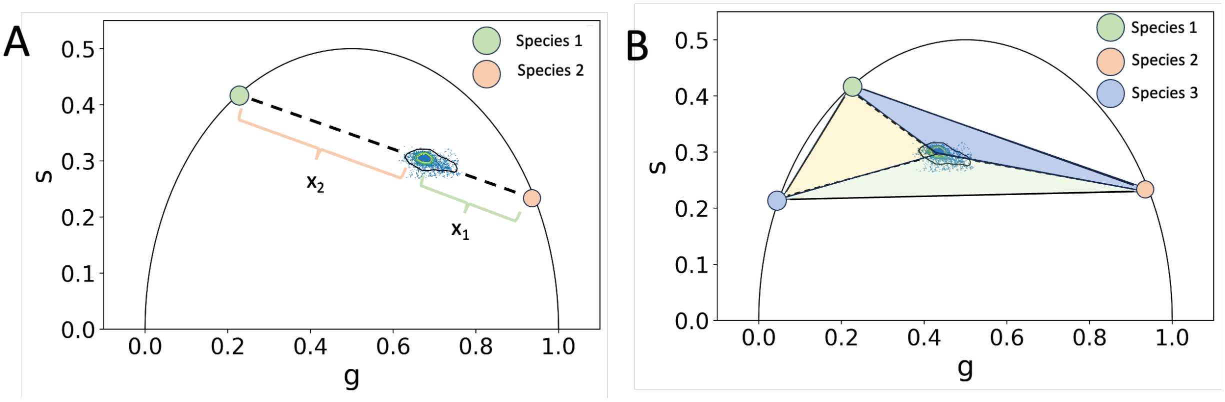


**Figure S1. Schematic representation of the graphical approach for assessing intensity fractions of coexisting species. A)** Linear approach for 2 coexisting species**. B)** Approach for 3 coexisting species.

| Sample | G | G std | S | S std | lifetime | lifetime std | sampled points | Contour PCA variability ratio |
|---|---|---|---|---|---|---|---|---|
| **irinotecan pH3.6** | 0.245019 | 0.007738 | 0.423314 | 0.006534 | 3.547728 | 0.010943 | 182338 | 0.581715 |
| **irinotecan pH7.2** | 0.238661 | 0.006967 | 0.421255 | 0.005676 | 3.595292 | 0.009852 | 170887 | 0.593733 |
| **irinotecan pH8.4** | 0.202396 | 0.006933 | 0.394718 | 0.005882 | 4.019535 | 0.009805 | 154348 | 0.579785 |
| **SN-38 pH4.25** | 0.282524 | 0.007482 | 0.403137 | 0.006235 | 3.517673 | 0.010582 | 244240 | 0.596726 |
| **SN-38 pH7.2** | 0.259005 | 0.008161 | 0.406107 | 0.006879 | 3.619584 | 0.011541 | 296999 | 0.589467 |

**Table S1. Comprehensive overview of the phasor measurements of Figure 3.** The data table shows the diverse impact of physical, chemical and biological changes on irinotecan, revealing alterations in the FLIM signal.

| pH | SN-38 lifetime [ns] | Irinotecan lifetime [ns] |
|---|---|---|
| **2.0** | 1.98 | 3.65 |
| **2.8** | 2.97 | 3.61 |
| **3.6** | 3.44 | 3.54 |
| **4.25** | 3.50 | 3.55 |
| **6.25** | 3.52 | 3.56 |
| **7.25** | 3.59 | 3.57 |
| **8.40** | 3.89 | 4.01 |
| **9.40** | 4.00 | 3.99 |
| **12.0** | 4.21 | 4.02 |

**Table S2. Comprehensive overview of the lifetime measurements of SN-38 and Irinotecan in PBS over the wide range of pH values under study, from 2 to 12.** The data table shows the diverse impact of pH changes on the lifetime of SN-38 and Irinotecan, revealing alterations in fluorescence behaviour.

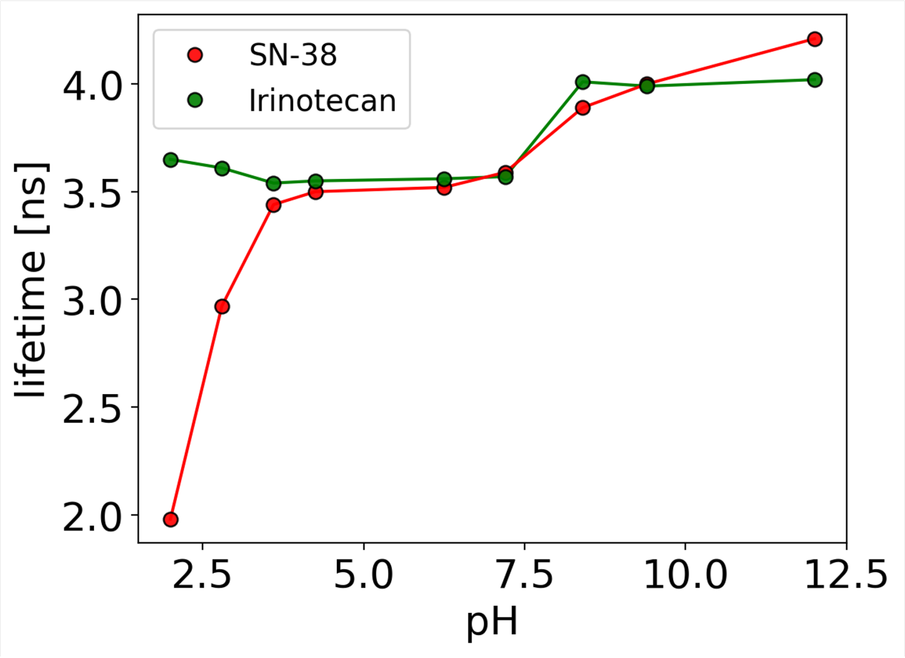


**Figure S2. Comparison of the fluorescence lifetime of irinotecan (green) and SN-38 (red) at pH values between 2 and 12.** Irinotecan exhibits minimal pH sensitivity compared to its metabolite.

| Distribution Parameter | Control | Cytokines |
|---|---|---|
| 25th Percentile | 2.4139 | 2.6355 |
| 50th Percentile | 2.5383 | 2.7703 |
| 75th Percentile | 2.669 | 2.9417 |
| Central tendency average | 2.5386 | 2.8161 |
| Spread (Standard Deviation) | 0.1921 | 0.2540 |
| Skewness | -0.0434 | 0.9799 |
| Mann Withney p-value | 0.0 | |
| Kolmogorov Smirnov p-value | 1.3746e-86 | |

**Table S3. Lifetime distribution data statistical comparison of INS1E cells (control) and Cytokines.** Statistical analysis of the lifetime distributions of Figure 4.

| Sample | ε [$M^{-1}$ $cm^{-1}$] | QY % |
|---|---|---|
| Crystal doxorubicin | 7510 ± 490 | 0.150 ± 0.004 |
| Membrane-bound doxorubicin | 10 340± 35 | 19.17 ± 0.4 |
| Free-in-solution doxorubicin | 10 340± 35 | 4.23 ± 0.09 |

**Table S4. Quantum yield and molar extinction coefficient of doxorubicin in distinct physical states: crystal, membrane-bound, and free in solution.** These values were sourced from reference [15] and were determined at the specific excitation wavelength of 488 nm.
